## Supplementary material for "Regional activity within the human amygdala varies with season, mood and illuminance": Online supplementary tables

**Short Title: Mood, season, light and the amygdala**

[1]GIGA-CRC-Human Imaging, University of Liège, Liège, 4000 Belgium
* shared first authorship

****Corresponding author**: Gilles Vandewalle, GIGA-Cyclotron Research Centre-In Vivo Imaging, Bâtiment B30, 8 Allée du Six Août, University of Liège-Sart Tilman, 4000 Liège, Belgium.

**Supplementary Table S1.** Characteristics of the participants included in the analyses.

| | **Mean (SD)** |
|---|---|
| **Number of Participants** | 29 |
| **Age** | 24y ± 3.1 |
| **Sex (M)** | 11 |
| **Mood (BDI-II)** | 7.9 ± 7.0 |
| **Number of participants per month (January to December)** | j:1; f:1; m:3; a:4; m:4; j:2; j:1; a:3; s:1; o:2; n:5; d:2 |
| **Anxiety (BAI)** | 5.1 ± 4.2 |
| **Sleep quality (PSQI)** | 4.1 ± 2.6 |
| **Seasonality (SPAQ)** | 1.2 ± 0.9 |
| **Chronotype (HO)** | 48.3 ± 7.6 |
| **Daytime sleepiness (ESS)** | 6.2 ± 2.9 |
| **Years of Education** | 14.2 ± 3.1 |
| **Sleep duration (night before fMRI protocol)** | 7.9 ± 0.7 |

BDI-II, Beck Depression Inventory-II [1]; BAI, Beck Anxiety Inventory [2]; PSQI, Pittsburgh Sleep Quality Index [3]; SPAQ, Seasonal Pattern Assessment Questionnaire [4]; HO, Horne and Östberg [5]; ESS, Epworth Sleepiness Scale [6].

**Supplementary Table 2.** Characteristics of the four light conditions used in fMRI protocol

| | Low BEL | Mid BEL | High BEL | Monochromatic light (589nm) |
|---|---|---|---|---|
| **Lux** | 47 | 116 | 240 | 7.5 |
| **Peak Spectral Irradiance (nm)** | 460 | 460 | 460 | 590 |
| **Melanopic EDI (lux ; ipRGCs)** | 37 | 92 | 190 | 0.16 |
| **Rhodopic EDI (lux ; Rods)** | 39 | 97 | 201 | 0.94 |
| **Cyanopic EDI (lux ; S-cones)** | 32 | 79 | 163 | 0 |
| **Chloropic EDI (lux; M-cones)** | 44 | 110 | 227 | 5 |
| **Erythropic EDI (lux ; L-cones)** | 46 | 113 | 233 | 8 |
| **Irradiance (µW/cm²)** | 15 | 36 | 75 | 1.4 |
| **Photon flux(1/cm²/s)** | $4.12^{E+13}$ | $1.02^{E+14}$ | $2.10^{E+14}$ | $4.24^{E+12}$ |
| **Log Photon Flux ($log_{10}$ (1/cm²/s)** | 13.61 | 14.01 | 14.32 | 12.63 |
| **Narrowband peak** | - | - | - | 589 |
| **Narrowband FWHM** | - | - | - | 10 |

Blue enriched light (BEL) (low, mid, and high) and monochromatic light (589nm).

**Supplementary Table S3.** Impact of the emotional content to the auditory stimuli on amygdala subparts.

| Main GLMM | | | | |
|---|---|---|---|---|
| **Effect** | **DF** | **F** | **P** | **Partial $R^2$** |
| **Stimulus type** | 1, 27.74 | 9.93 | **<.0039** | **0.26** |
| **Amygdala subpart** | 9, 499.9 | 2.13 | **<.0256** | **0.04** |
| **Amygdala subparts x stimulus type** | 9, 499.9 | 0.56 | 0.85 | - |
| **Age** | 1, 24.98 | 2.20 | 0.15 | - |
| **Sex** | 1, 25 | 0.69 | 0.41 | - |
| **BMI** | 1, 24.97 | 1.69 | 0.20 | - |
| **Least Squares Means** | | | | |
| **Amygdala subpart** | **t Value** | **P-value** | | |
| 1 | 1.99 | **0.0494** | | |
| 2 | 0.71 | 0.4787 | | |
| 3 | 3.46 | **0.0008** | | |
| 4 | 1.75 | 0.0834 | | |
| 5 | 3.64 | **0.0005** | | |
| 6 | 1.59 | 0.1155 | | |
| 7 | 4.08 | **<.0001** | | |
| 8 | 2.92 | **0.0044** | | |
| 9 | 3.18 | **0.0020** | | |
| 10 | 2.17 | **0.0327** | | |

Amygdala subpart (refer to Fig. 2): (1) Lateral nucleus, (2) Intermediate and dorsal basolateral nucleus, (3) Basomedial nuclei, (4) Central nucleus, (5) Medial and Cortical nuclei, (6) Ventral basolateral nucleus and Paralaminar nucleus, (7) amygdala transition area composed of Amygdalocortical area, Amygdalohippocampal area, Periamygdaloid cortex, (8) Amygdalostriatal, (9) Anterior amygdaloid area, (10) Intercalated nuclei**.**

BMI: body mass index

**Supplementary Table S4.** Impact of season and affective state on the activity of amygdala subparts.

| a. GLMM | | | | |
|---|---|---|---|---|
| **Effect** | **DF** | **F** | **P** | **Partial $R^2$** |
| Stimulus type | 1, 23.85 | 8.85 | **0.0066** | 0.27 |
| Amygdala subpart | 6, 272.7 | 1.58 | 0.15 | - |
| Mood | 1, 17.81 | 17.46 | **0.0006** | 0.495 |
| Mood* Amygdala subpart | 6, 272.4 | 2.99 | **0.0076** | 0.062 |
| Time-of-year # | 1, 17.85 | 12.62 | **0.0023** | 0.41 |
| Season * Amygdala subpart | 6, 272.6 | 4.66 | **0.0002** | 0.09 |
| Mood* Time-of-year | 1, 17.79 | 12.18 | **0.0027** | 0.41 |
| Mood * Time-of-year * Amygdala subpart | 6, 272.4 | 4.03 | **0.0007** | 0.08 |
| Age | 1, 17.84 | 7.77 | **0.0122** | 0.3 |
| Sex | 1, 17.88 | 2.66 | 0.12 | - |
| BMI | 1, 17.82 | 1.37 | 0.26 | - |

b. **Time-of-year**

| **Amygdala subpart** | **T** | **P** | $P_{corrected}$ |
|---|---|---|---|
| 1 | 2.22 | 0.0287 | 0.1570 |
| 3 | 3.54 | 0.0006 | **0.0040** |
| 5 | 2.13 | 0.0356 | 0.1917 |
| 7 | 1.68 | 0.0963 | 0.4555 |
| 8 | -0.96 | 0.3384 | 0.9219 |
| 9 | 4.96 | <.0001 | **<.0001** |
| 10 | 2.06 | 0.0417 | 0.2211 |

c. **Time-of-year * amygdala subpart**

| **Amygdala subpart contrast** | **T** | **P** | **$P_{corrected}$** |
|---|---|---|---|
| 1 vs. 3 | -1.11 | 0.2665 | 0.9261 |
| 1 vs. 5 | 0.08 | 0.9401 | 1.0000 |
| 1 vs. 7 | 0.42 | 0.6720 | 0.9992 |
| 1 vs. 8 | 2.68 | **0.0078** | 0.1073 |
| 1 vs. 9 | -2.32 | **0.0210** | 0.2348 |
| 1 vs. 10 | 0.13 | 0.8951 | 1.0000 |
| 3 vs. 5 | 1.19 | 0.2356 | 0.9000 |
| 3 vs. 7 | 1.52 | 0.1291 | 0.7305 |
| 3 vs. 8 | 3.80 | **0.0002** | **0.0037** |
| 3 vs. 9 | -1.21 | 0.2268 | 0.8925 |
| 3 vs. 10 | 1.25 | 0.2141 | 0.8796 |
| 5 vs. 7 | 0.35 | 0.7269 | 0.9998 |
| 5 vs. 8 | 2.61 | 0.0096 | 0.1269 |
| 5 vs. 9 | -2.40 | **0.0172** | 0.2021 |
| 5 vs. 10 | 0.06 | 0.9548 | 1.0000 |
| 7 vs. 8 | 2.22 | **0.0271** | 0.2817 |
| 7 vs. 9 | -2.71 | **0.0071** | 0.0992 |

| | | | |
|---|---|---|---|
| 7 vs. 10 | -0.29 | 0.7692 | 0.9998 |
| 8 vs. 9 | -5.00 | **<.0001** | **<.0001** |
| 8 vs. 10 | -2.55 | **0.0113** | 0.1457 |
| 9 vs. 7 | 2.45 | **0.0148** | 0.1806 |
| **d. <u>Mood</u>** | | | |
| **Amygdala subpart** | **T** | **P** | **$P_{corrected}$** |
| 1 | 1.85 | 0.0668 | 0.3350 |
| 3 | 2.17 | **0.0328** | 0.1764 |
| 5 | 4.28 | **<.0001** | **0.0002** |
| 7 | 4.64 | **<.0001** | **<.0001** |
| 8 | 0.40 | 0.6881 | 0.9994 |
| 9 | 2.60 | **0.0107** | 0.0614 |
| 10 | 2.46 | **0.0156** | 0.0879 |
| **e. <u>Mood * amygdala subpart</u>** | | | |
| **Amygdala subpart contrast** | **T** | **P** | **$P_{corrected}$** |
| 1 vs. 3 | -0.26 | 0.7926 | 0.9999 |
| 1 vs. 5 | -2.05 | **0.0416** | 0.3814 |
| 1 vs. 7 | -2.36 | **0.0190** | 0.2163 |
| 1 vs. 8 | 1.22 | 0.2224 | 0.8823 |
| 1 vs. 9 | -0.64 | 0.5243 | 0.9940 |
| 1 vs. 10 | -0.51 | 0.6086 | 0.9979 |
| 3 vs. 5 | -1.78 | 0.0755 | 0.5583 |
| 3 vs. 7 | -2.10 | **0.0370** | 0.3503 |
| 3 vs. 8 | 1.49 | 0.1384 | 0.7519 |
| 3 vs. 9 | -0.38 | 0.7079 | 0.9997 |
| 3 vs. 10 | -0.25 | 0.8032 | 0.9999 |
| 5 vs. 7 | -0.32 | 0.7518 | 0.9998 |
| 5 vs. 8 | 3.27 | **0.0012** | 0.0198 |
| 5 vs. 9 | 1.40 | 0.1613 | 0.7987 |
| 5 vs. 10 | 1.53 | 0.1261 | 0.7218 |
| 7 vs. 8 | 3.58 | **0.0004** | **0.0083** |
| 7 vs. 9 | 1.72 | 0.0871 | 0.6051 |
| 7 vs. 10 | 1.85 | 0.0658 | 0.5123 |
| 8 vs. 9 | -1.86 | 0.0643 | 0.5051 |
| 8 vs. 10 | -1.74 | 0.0838 | 0.5925 |
| 9 vs. 7 | 0.13 | 0.8997 | 1.0000 |

Amygdala subparts (refer to Fig. 2): (1) Lateral nucleus, (3) Basomedial nuclei, (5) Medial and Cortical nuclei, (7) amygdala transition area composed of Amygdalocortical area, Amygdalohippocampal area, Periamygdaloid cortex, (8) Amygdalostriatal, (9) Anterior amygdaloid area, (10) Intercalated nuclei**.**

BMI: body mass index

**Supplementary Table S5.** Impact of light illuminance on the activity of amygdala subparts.

| **GLMM** | | | | |
|---|---|---|---|---|
| **Effect** | **DF** | **F** | **P** | **Partial $R^2$** |
| Stimulus type | 1, 27.24 | 6.32 | **0.0181** | 0.19 |
| Amygdala subpart | 6, 331.4 | 7.05 | **<.0001** | 0.11 |
| Stimulus type * Amygdala subpart | 6, 331.4 | 1.42 | 0.21 | - |
| Age | 1, 25.06 | 1.23 | 0.28 | - |
| Sex | 1, 25.05 | 1.41 | 0.25 | - |
| BMI | 1, 25.12 | 0.11 | 0.74 | - |
| **Illuminance - Least Squares Means** | | | | |
| **Amygdala subpart** | **T** | **P** | | |
| 1 | -1.90 | 0.0623 | | |
| 3 | -4.12 | **0.0001** | | |
| 5 | -4.47 | **<.0001** | | |
| 7 | -3.68 | **0.0005** | | |
| 8 | -0.30 | 0.7675 | | |
| 9 | -3.92 | **0.0002** | | |
| 10 | -0.63 | 0.5283 | | |
| **Illuminance * amygdala subpart** | | | | |
| **Amygdala subpart contrast** | **T** | **P** | **$P_{corrected}$** | |
| 1 vs. 3 | 2.39 | 0.0172 | 0.2087 | |
| 1 vs. 5 | 2.77 | 0.0059 | 0.0893 | |
| 1 vs. 7 | 1.92 | 0.0561 | 0.4693 | |
| 1 vs. 8 | -1.72 | 0.0873 | 0.6069 | |
| 1 vs. 9 | 2.2 | 0.0283 | 0.2978 | |
| 1 vs. 10 | -1.36 | 0.1741 | 0.8205 | |
| 3 vs. 5 | 0.38 | 0.7072 | 0.9998 | |
| 3 vs. 7 | -0.48 | 0.6329 | 0.9993 | |
| 3 vs. 8 | -4.1 | <.0001 | 0.0005 | |
| 3 vs. 9 | -0.16 | 0.8765 | 1 | |
| 3 vs. 10 | -3.76 | 0.0002 | 0.0025 | |
| 5 vs. 7 | -0.85 | 0.3937 | 0.98 | |
| 5 vs. 8 | -4.47 | <.0001 | 0.0002 | |
| 5 vs. 9 | -0.53 | 0.5995 | 0.9988 | |
| 5 vs. 10 | -4.13 | <.0001 | 0.0005 | |
| 7 vs. 8 | -3.62 | 0.0003 | 0.005 | |
| 7 vs. 9 | 0.32 | 0.7529 | 0.9998 | |
| 7 vs. 10 | -3.28 | 0.0012 | 0.0206 | |
| 8 vs. 9 | 3.89 | 0.0001 | 0.0011 | |
| 8 vs. 10 | 0.36 | 0.7191 | 0.9998 | |
| 9 vs. 7 | -3.54 | 0.0005 | 0.0074 | |

Amygdala subpart (refer to Fig. 2): (1) Lateral nucleus, (3) Basomedial nuclei, (5) Medial and Cortical nuclei, (7) amygdala transition area composed of Amygdalocortical area, Amygdalohippocampal area, Periamygdaloid cortex, (8) Amygdalostriatal, (9) Anterior amygdaloid area, (10) Intercalated nuclei**.** BMI: body mass index

**Supplementary Table S6.** Impact of each illuminance on the activity of the amygdala subparts affected by light exposure.

| GLMM | | | | |
|---|---|---|---|---|
| **Effect** | **DF** | **F** | **P** | **Partial $R^2$** |
| Stimulus type | 1, 244.1 | 9.12 | **0.0028** | 0.04 |
| Illuminance | 4, 244.1 | 6.09 | **0.0001** | 0.09 |
| Illuminance * Stimulus type | 4, 244.2 | 1.65 | 0.1617 | - |
| Amygdala subpart | 3, 813.4 | 0.1 | 0.9587 | - |
| Stimulus type * Amygdala subpart | 3, 813.6 | 3.08 | **0.0269** | 0.01 |
| Illuminance * Amygdala subpart | 12, 813.5 | 0.48 | 0.9294 | - |
| Stimulus type * Illuminance * Amygdala subpart | 12, 813.5 | 0.26 | 0.9942 | - |
| Age | 1, 24.98 | 2.9 | 0.1008 | - |
| Sex | 1, 24.99 | 0.3 | 0.5885 | - |
| BMI | 1, 24.98 | 1.84 | 0.187 | - |

**Illuminance per amygdala subpart**

| **Amygdala subpart** | Contrast | **T** | **P** | $P_{corrected}$ |
|---|---|---|---|---|
| 3 | 0 vs. 0.16 | -0.59 | 0.55 | 0.97 |
| 3 | 0 vs. 37 | 1.01 | 0.31 | 0.85 |
| 3 | 0 vs. 92 | -0.07 | 0.94 | 1 |
| 3 | 0 vs. 190 | 2.76 | **0.0059** | **0.046** |
| 3 | 0.16 vs. 37 | 1.6 | 0.11 | 0.5 |
| 3 | 0.16 vs. 92 | 0.51 | 0.61 | 0.97 |
| 3 | 0.16 vs. 190 | 3.35 | **0.0008** | **0.0075** |
| 3 | 37 vs. 92 | -1.09 | 0.28 | 0.81 |
| 3 | 37 vs. 190 | 1.74 | 0.083 | 0.41 |
| 3 | 92 vs. 190 | 2.84 | **0.0047** | **0.0375** |
| 5 | 0 vs. 0.16 | -0.07 | 0.94 | 1 |
| 5 | 0 vs. 37 | 0.69 | 0.49 | 0.96 |
| 5 | 0 vs. 92 | 0.35 | 0.72 | 0.99 |
| 5 | 0 vs. 190 | 3.07 | **0.0022** | **0.019** |
| 5 | 0.16 vs. 37 | 0.76 | 0.45 | 0.94 |
| 5 | 0.16 vs. 92 | 0.42 | 0.67 | 0.99 |
| 5 | 0.16 vs. 190 | 3.14 | **0.0018** | **0.015** |
| 5 | 37 vs. 92 | -0.33 | 0.74 | 0.99 |
| 5 | 37 vs. 190 | 2.38 | **0.0175** | 0.12 |
| 5 | 92 vs. 190 | 2.69 | **0.0072** | 0.056 |
| 7 | 0 vs. 0.16 | 0.19 | 0.85 | 0.99 |
| 7 | 0 vs. 37 | 1.01 | 0.31 | 0.85 |
| 7 | 0 vs. 92 | 0.34 | 0.74 | 0.99 |
| 7 | 0 vs. 190 | 2.53 | **0.0116** | 0.085 |
| 7 | 0.16 vs. 37 | 0.82 | 0.41 | 0.93 |
| 7 | 0.16 vs. 92 | 0.15 | 0.88 | 0.99 |
| 7 | 0.16 vs. 190 | 2.34 | **0.0195** | 0.13 |

| | | | | |
|---|---|---|---|---|
| 7 | 37 vs. 92 | -0.67 | 0.50 | 0.96 |
| 7 | 37 vs. 190 | 1.51 | 0.13 | 0.55 |
| 7 | 92 vs. 190 | 2.19 | **0.029** | 0.18 |
| 9 | 0 vs. 0.16 | 0.28 | 0.78 | 0.99 |
| 9 | 0 vs. 37 | -0.04 | 0.97 | 1 |
| 9 | 0 vs. 92 | 0.62 | 0.54 | 0.97 |
| 9 | 0 vs. 190 | 3.55 | **0.0004** | **0.0038** |
| 9 | 0.16 vs. 37 | -0.32 | 0.75 | 0.99 |
| 9 | 0.16 vs. 92 | 0.34 | 0.73 | 0.99 |
| 9 | 0.16 vs. 190 | 3.23 | **0.0013** | **0.011** |
| 9 | 37 vs. 92 | 0.66 | 0.51 | 0.97 |
| 9 | 37 vs. 190 | 3.58 | **0.0004** | **0.0033** |
| 9 | 92 vs. 190 | 2.91 | **0.0038** | **0.031** |

Activity of the 4 subparts showing a significant impact of overall illuminance changes (cf. Table S5 and Fig. 3C-D) was extracted from a separate analysis of the fMRI data which considered separately each illuminance (cf. methods). As in the main analysis, the GLMM including these activity estimates as dependent variable confirmed that there was a significant impact of stimulus type and light level. The 4 subparts did not differ between each other and their changes in responses with illuminance change were not different, which is not surprising given the fact that the four subparts were selected based on their overall response to illuminance change.

Post hoc contrasts indicate that the activity of the four subparts significantly decreased under the highest illuminance (190 mel EDI) when compared to darkness (0 mel EDI lux) and the lowest illuminance level (.16 mel EDI lux) and with 37 mel EDI illuminance in some cases (though significance was not surviving correction for multiple comparisons in some cases). Together with the visual inspection of Figure 3C, the overall impression is of linear decrease in activity with increasing illuminance (in line with the output of the analyses considering illuminance variation as a whole – cf. Table S5).

Amygdala subpart (refer to Fig. 2): (3) Basomedial nuclei, (5) Medial and Cortical nuclei, (7) amygdala transition area composed of Amygdalocortical area, Amygdalohippocampal area, Periamygdaloid cortex, (9) Anterior amygdaloid area**.**

BMI: body mass index

**Supplementary Table S7.** Impact of season and affective state on the impact of light exposure on the activity of amygdala subparts.

| GLMM | | | | |
|---|---|---|---|---|
| **Effect** | **DF** | **F** | **P** | **Partial $R^2$** |
| Stimulus type | 1, 23.13 | 9.01 | **0.0063** | 0.28 |
| Amygdala subpart | 6, 272 | 5.63 | **<.0001** | 0.11 |
| Mood | 1, 18 | 1.23 | 0.28 | - |
| Mood * Amygdala subpart | 6, 272 | 2.24 | **0.04** | 0.05 |
| Time-of-year [#] | 1, 18 | 0.02 | 0.89 | - |
| **Time-of-year* Amygdala subpart** | 6, 272 | 2.94 | **0.0085** | 0.06 |
| Mood * Season | 1, 18.01 | 0.25 | 0.62 | - |
| Mood * Time-of-year * Amygdala subpart | 6, 272 | 0.87 | 0.52 | - |
| Age | 1, 18.07 | 0.97 | 0.34 | - |
| Sex | 1, 18.02 | 0.6 | 0.45 | - |
| BMI | 1, 18.05 | 0.09 | 0.77 | - |

**<u>Time-of-year</u>**

| **Amygdala subpart** | **T** | **P** | $P_{corrected}$ |
|---|---|---|---|
| 1 | 0.58 | 0.56 | 0.98 |
| 3 | 0.27 | 0.79 | 0.99 |
| 5 | 1.06 | 0.29 | 0.76 |
| 7 | -1.71 | **0.096** | 0.32 |
| 8 | -0.92 | 0.36 | 0.85 |
| 9 | 1.14 | 0.26 | 0.70 |
| 10 | 0.36 | 0.72 | 0.99 |

**<u>Time-of-year * amygdala subpart</u>**

| **Amygdala subpart contrast** | **T** | **P** | **$P_{corrected}$** |
|---|---|---|---|
| 1 vs. 3 | 0.36 | 0.72 | 0.99 |
| 1 vs. 5 | -0.56 | 0.58 | 0.99 |
| 1 vs. 7 | 2.64 | 0.0089 | 0.12 |
| 1 vs. 8 | 1.73 | 0.085 | 0.6 |
| 1 vs. 9 | -0.65 | 0.52 | 0.99 |
| 1 vs. 10 | 0.26 | 0.8 | 0.99 |
| 3 vs. 5 | -0.92 | 0.36 | 0.97 |
| 3 vs. 7 | 2.27 | **0.0237** | 0.26 |
| 3 vs. 8 | 1.37 | 0.17 | 0.83 |
| 3 vs. 9 | -1.01 | 0.31 | 0.95 |
| 3 vs. 10 | -0.11 | 0.92 | 1 |
| 5 vs. 7 | 3.19 | **0.0016** | **0.0244** |
| 5 vs. 8 | 2.29 | **0.023** | 0.26 |
| 5 vs. 9 | -0.09 | 0.93 | 1 |
| 5 vs. 10 | 0.81 | 0.42 | 0.98 |
| 7 vs. 8 | -0.9 | 0.37 | 0.97 |

| | | | |
|---|---|---|---|
| 7 vs. 9 | -3.28 | **0.0012** | 0.0185 |
| 7 vs. 10 | -2.38 | **0.018** | 0.2167 |
| 8 vs. 9 | -2.38 | **0.0182** | 0.218 |
| 8 vs. 10 | -1.48 | 0.1413 | 0.7691 |
| 9 vs. 7 | 0.91 | 0.3662 | 0.9732 |
| **Mood** | | | |
| **Amygdala subpart contrast** | **T** | **P** | **$P_{corrected}$** |
| 1 | 0.66 | 0.51 | 0.96 |
| 3 | 1.73 | 0.091 | 0.29 |
| 5 | 1.53 | 0.14 | 0.41 |
| 7 | 0 | 0.99 | 1 |
| 8 | -0.21 | 0.84 | 1 |
| 9 | 2.21 | **0.033** | 0.12 |
| 10 | 0.47 | 0.64 | 0.99 |
| **Mood * amygdala subpart** | | | |
| 1 vs. 3 | -1.23 | 0.22 | 0.87 |
| 1 vs. 5 | -0.99 | 0.32 | 0.95 |
| 1 vs. 7 | 0.77 | 0.44 | 0.99 |
| 1 vs. 8 | 1 | 0.32 | 0.95 |
| 1 vs. 9 | -1.78 | 0.077 | 0.56 |
| 1 vs. 10 | 0.23 | 0.819 | 1 |
| 3 vs. 5 | 0.24 | 0.81 | 1 |
| 3 vs. 7 | 2 | **0.046** | 0.42 |
| 3 vs. 8 | 2.23 | **0.027** | 0.29 |
| 3 vs. 9 | -0.55 | 0.58 | 0.99 |
| 3 vs. 10 | 1.46 | 0.15 | 0.76 |
| 5 vs. 7 | 1.76 | 0.079 | 0.57 |
| 5 vs. 8 | 1.99 | **0.0475** | 0.42 |
| 5 vs. 9 | -0.79 | 0.43 | 0.99 |
| 5 vs. 10 | 1.22 | 0.22 | 0.88 |
| 7 vs. 8 | 0.24 | 0.81 | 1 |
| 7 vs. 9 | -2.54 | **0.0116** | 0.15 |
| 7 vs. 10 | -0.54 | 0.58 | 0.99 |
| 8 vs. 9 | -2.77 | **0.006** | 0.084 |
| 8 vs. 10 | -0.77 | 0.44 | 0.99 |
| 9 vs. 7 | 2 | **0.046** | 0.41 |

Amygdala subparts (refer to Fig. 2): (1) Lateral nucleus, (3) Basomedial nuclei, (5) Medial and Cortical nuclei, (7) amygdala transition area composed of Amygdalocortical area, Amygdalohippocampal area, Periamygdaloid cortex, (8) Amygdalostriatal, (9) Anterior amygdaloid area, (10) Intercalated nuclei**.**

BMI: body mass index
